# Reliable individual differences in fine-grained cortical functional architecture

**DOI:** 10.1101/296012

**Authors:** Ma Feilong, Samuel A. Nastase, J. Swaroop Guntupalli, James V. Haxby

## Abstract

Fine-grained functional organization of cortex is not well-conserved across individuals. As a result, individual differences in cortical functional architecture are confounded by topographic idiosyncrasies—i.e., differences in functional-anatomical correspondence. In this study, we used hyperalignment to align information encoded in topographically variable patterns to study individual differences in fine-grained cortical functional architecture in a common representational space. We characterized the structure of individual differences using three common functional indices, and assessed the reliability of this structure across independent samples of data in a natural vision paradigm. Hyperalignment markedly improved the reliability of individual differences across all three indices by resolving topographic idiosyncrasies and accommodating information encoded in spatially fine-grained response patterns. Our results demonstrate that substantial individual differences in cortical functional architecture exist at fine spatial scales, but are inaccessible with anatomical normalization alone.

## 1 Introduction

Functional architecture of the human brain is relatively consistent across individuals at a coarse scale, but idiosyncrasies in functional topography become increasingly apparent at finer scales. Large-scale brain systems can be reliably identified across individuals by functional connectivity (Power et al., 2011; Yeo et al., 2011), but there is profound interindividual variability in system details (Gordon et al., 2017a, 2017b). At the areal level, category-selective regions can be localized to anatomical landmarks (Weiner et al., 2018, 2014), though the locus can differ across individuals by millimeters or centimeters, along with variability in size and shape (Zhen et al., 2017, 2015). Furthermore, within a brain area, between-subject classification of response patterns is typically considerably worse than within-subject classification (e.g., Cox and Savoy, 2003), indicating that fine-grained functional architecture is not well-aligned macroanatomically. With state-of-the-art cortical surface-based alignment (Fischl, 2012), the mismatch between brain function and anatomy can be reduced but not eliminated (Duncan et al., 2009; Frost and Goebel, 2012; Weiner et al., 2018). Therefore, it is problematic to assume that a given anatomical location or topographic conformation will have the same functional role across brains. Individual differences in neural function are confounded by individual differences in functional-anatomical correspondence.

Hyperalignment (Guntupalli et al., 2016, in press; Haxby et al., 2011) is a family of methods that can disentangle functional variability from anatomical variability. Hyperalignment projects features (voxels or surface vertices) from a brain to a common high-dimensional space through linear transformations. In this common space, the same features from different individuals will share similar functional properties instead of the same anatomical locations or topographic conformations. Hyperalignment decomposes the original fMRI data of each individual into two parts: a transformation matrix, which reflects topographic properties of the individual’s functional activations; and a new data matrix in the common space, which reflects shared, stimulus-driven responses. This hyperaligned data matrix provides an opportunity to study brain functions without confounds from topographic variability.

Besides separating interindividual variability in brain function from that in functional topography, hyperalignment also makes efficient use of functional neuroimaging data by utilizing spatially fine-grained pattern information. For example, between-subject classification improves severalfold by using hyperalignment instead of anatomical alignment (Guntupalli et al., in press, 2016; Haxby et al., 2011). Fine-grained information exists across all of cortex (Guntupalli et al., in press, 2016; Haxby et al., 2014, 2011), but cannot be modeled across individuals using anatomical alignment alone. Many researchers opt to sacrifice fine-grained information to boost coarse-grained information through spatial smoothing or averaging (Carp, 2012), making fine-grained information inaccessible to further analysis.

Recent advances have increased the utility of fMRI for studying the neural bases of individual differences (Dubois and Adolphs, 2016). Functional responses and connectivity measured by fMRI can predict an individual’s intelligence (Heuvel et al., 2009; Smith et al., 2013), creativity (Beaty et al., 2018), personality (Adelstein et al., 2011; DeYoung et al., 2010), and can facilitate diagnosis (Arbabshirani et al., 2017; Wolfers et al., 2015) and prognosis (Gabrieli et al., 2015) of psychiatric disorders. However, it is still common practice to use feature sets derived by averaging voxels within a region or a network (Wolfers et al., 2015).

In this study, we used hyperalignment to examine individual differences in cortical functional architecture in a common representational space. Critically, this common space resolves idiosyncrasies in functional-anatomical correspondence. We indexed cortical functional architecture based on response profiles, functional connectivity, and representational geometry, and assessed the reliability of individual differences across independent samples of data using a natural vision paradigm. We found that with hyperaligned data, individual differences in cortical functional architecture were more reliable, and this increased reliability results primarily from the incorporation of fine-scale cortical functional architecture into our model. Our results suggest that substantial individual differences exist at a fine spatial scale, but are obscured by idiosyncratic functional-anatomical correspondence; hyperalignment can reveal these fine-scale individual differences, and make observed individual differences in cortical functional architecture more reliable.

## 2 Materials and methods

### 2.1 Participants

Twenty healthy young adults (mean age ± standard deviation: 24.4 ± 3.4 years, 12 females) participated in this study. All participants were right-handed, with normal hearing and normal or corrected-to-normal vision, and no known history of neurological illness. They gave written, informed consent, and were paid for their participation. The study was approved by the Institutional Review Board of Dartmouth College.

### 2.2 Stimuli and design

Participants watched a full-length audiovisual movie, *Raiders of the Lost Ark*, while fMRI data were collected. The movie was divided into eight parts, each 14-15 minutes in duration. Participants viewed four parts of the movie in each of the two scanning sessions, and were taken out of the scanner between the two sessions for a break.

The video was projected from an LCD projector onto a rear projection screen, and then reflected through a mirror on the head coil. The corresponding visual angles subtended approximately 22.7° horizontally and 17° vertically. The audio was played through MR-compatible headphones (MR confon GmbH, Magdeburg, Germany). Participants were instructed to pay attention to the movie and enjoy.

### 2.3 MRI acquisition

MR images were acquired using a 3T Philips Intera Achieva scanner with a 32-channel head coil at the Dartmouth Brain Imaging Center.

Functional images comprised 80 × 80 × 42 3 mm isotropic voxels, providing whole brain coverage. They were acquired every 2.5 seconds with an echo planar imaging (EPI) sequence (TR = 2.5 s, TE = 35 ms, flip angle = 90°, 80 × 80 matrix, FOV = 240 mm × 240 mm, SENSE reduction factor = 2, 42 interleaved axial slices). The length of a run was adjusted to match the length of the corresponding movie part, consisting of 326 to 344 volumes each. In total, we acquired 2718 functional images per participant during 8 runs of movie watching (approximately 2 hrs of functional data per participant).

A high resolution T1-weighted image (0.9375 mm × 0.9375 mm × 1.0 mm voxel resolution) was also acquired in each session with an MPRAGE sequence (TR = 8.2 ms, TE = 3.7 ms, flip angle = 8°, 256 × 256 matrix, FOV = 240 mm × 240 mm, 220 axial slices), except for one session of one participant.

### 2.4 MRI preprocessing

MRI data were first preprocessed using the fmriprep software version 1.0.0-rc2 (https://github.com/poldracklab/fmriprep). T1-weighted images were corrected for bias field (Tustison et al., 2010) and skullstripped using antsBrainExtraction.sh. High resolution cortical surfaces were reconstructed with FreeSurfer (Fischl, 2012; http://surfer.nmr.mgh.harvard.edu/) using all available anatomical images, and registered to the fsaverage template (Fischl et al., 1999). Functional data were motion corrected using MCFLIRT (Jenkinson et al., 2002), and resampled to the fsaverage template based on boundary-based registration (Greve and Fischl, 2009). After these steps, functional data from all participants were in alignment with the fsaverage template based on cortical folding patterns.

Further preprocessing steps were performed using Python scripts based on PyMVPA (Hanke et al., 2009; http://www.pymvpa.org/). First, functional data were further downsampled to a standard cortical surface mesh with 18,742 vertices across both hemispheres (3 mm vertex spacing; 20,484 vertices before removing non-cortical vertices), and data acquired during overlapping movie segments were discarded (8 TRs, 20 seconds, for each of runs 2-8). Then, nuisance regressors—6 motion parameters and their derivatives, global signal, framewise displacement (Power et al., 2014), 6 principal components from cerebrospinal fluid and white matter (aCompCor; Behzadi et al., 2007), and up to second order polynomial trends—were partialed out from functional data separately for each run. Finally, the residual time series of each surface vertex in each run was normalized to zero mean and unit variance.

### 2.5 Hyperalignment

First, we created a common representational space to hyperalign functional data based on an independent dataset. The dataset comprised responses to the same movie from 11 different participants (Guntupalli et al., 2016; Haxby et al., 2011). We preprocessed the 11 participants’ data through the same pipeline, and hyperaligned responses to the entire movie for all subjects using searchlight hyperalignment (Guntupalli et al., 2016) with a 20-mm searchlight radius. For a given searchlight, the hyperalignment algorithm uses the Procrustes transformation to rotate each subject’s feature space so as to best align response trajectories across individuals. The local transformations for each searchlight are then aggregated, forming a single, sparse transformation matrix for each cortical hemisphere. The hyperaligned data of the 11 participants were then averaged and normalized to unit variance to serve as the final common representational space.

Then, we derived hyperalignment transformations for each of the 20 participants to the common representational space based on their responses to the first half of the movie (runs 1-4; different from the 11 participants), and applied these transformations to data from the second half of the movie (runs 5-8). The following analyses of individual differences were based on the second half of the movie for these 20 participants, and are thus completely independent of the data used for deriving the common space and hyperalignment parameter estimation. Note that all data were anatomically aligned according to sulcal curvature on the cortical surface prior to hyperalignment (see 2.4).

### 2.6 Measuring the reliability of individual differences

We used surface-based searchlight analysis (Kriegeskorte et al., 2006; Oosterhof et al., 2011) to measure individual differences in local cortical functional architecture. Within each searchlight (9 mm radius), we modeled individual differences in local cortical functional architecture as a 20 subjects × 20 subjects similarity matrix, which we refer to as an individual differences matrix (IDM). Each entry in an IDM is the similarity between functional indices (see 2.7) for a pair of individuals; i.e., the Pearson correlation between two vectors, one for each individual, capturing a multivariate neural signature based on a given functional index. By modeling the similarity structure (Kriegeskorte and Kievit, 2013) between individuals, IDMs can be compared across different stimuli (e.g., different parts of the movie data) and different functional indices.

We utilized this feature of IDMs and assessed the reliability of individual differences by comparing IDMs based on different parts of the movie data. Within each searchlight, we split responses to the second half of the movie into two parts (runs 5-6, 7-8), computed an IDM from each part, and evaluated the reliability of the individual differences structure by comparing the two matrices. See Fig. 1A for a schematic of this procedure. Because IDMs are symmetric and the diagonals are not informative, we quantified reliability as the Pearson correlation between the vectorized upper triangles of the IDMs. The reliability values from all searchlights form a reliability map of individual differences.

**Fig. 1.**
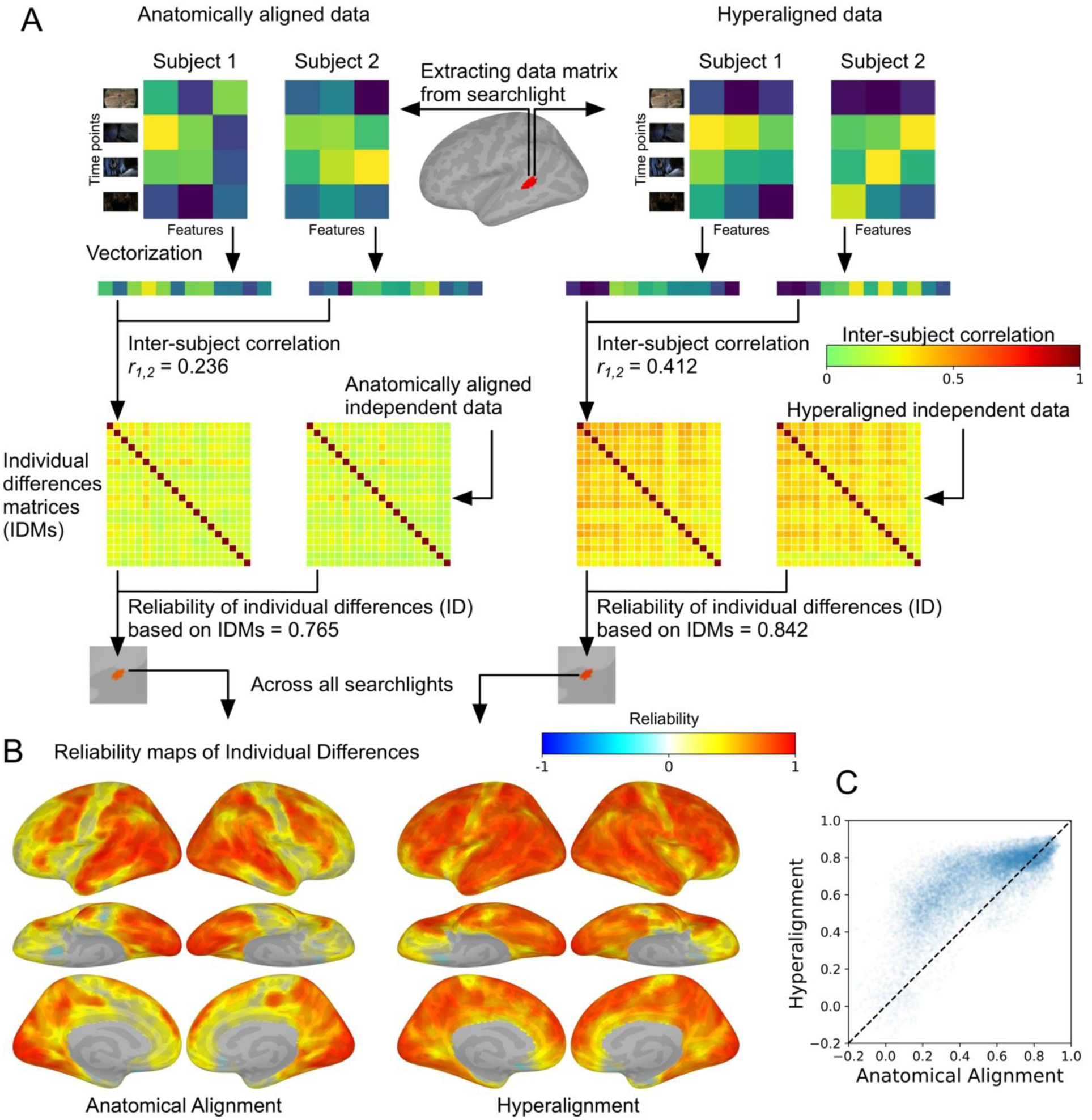
Measuring the reliability of individual differences in local response profiles. (A)Schematic illustration of the analysis pipeline. Using either anatomically aligned or hyperaligned data, within each searchlight, a time-points (i.e., TRs) by features (i.e., vertices) data matrix was extracted from each subject, and vectorized into a vector of response profiles. The similarity (i.e., correlation) of the vectors for each pair of subjects was computed, forming a subjects-by-subjects similarity matrix (individual differences matrix; IDM) capturing individual differences. Two IDMs were obtained based on independent data (responses to two different parts of the movie) from the same group of people, and the reliability of individual differences was measured as the correlation of their vectorized upper triangles. Note that we used a similar procedure to measure the reliability of individual differences in functional connectivity patterns and representational geometries (see text for details and Fig. 2). (B) Reliability maps of individual differences. These maps depict the searchlight reliabilities of individual differences in local response profiles based on anatomically aligned data (left) and hyperaligned data (right). (C) Scatterplot of searchlight reliabilities of IDMs in anatomically-aligned data (x-axis) and hyperaligned data (y-axis). In general, individual differences are more reliable with hyperaligned data. Most searchlights (82.1 %) showed an increase in the reliability of individual differences with hyperalignment, and the average reliability across all searchlights increased from 0.540 to 0.693.

Using each of the functional indices, we obtained two reliability maps of individual differences, one based on anatomically aligned data and the other based on hyperaligned data. To simplify the comparison between reliability maps, we first focused on comparing the average reliability difference across all searchlights. Then, we counted the percentage of searchlights that showed the same direction of effect as the average reliability. If the average reliability based on one alignment method is higher, and the reliabilities in most searchlights are also higher with the same alignment method, then the average reliability difference is not likely to be driven by outlier searchlights, and the average reliability difference is a good summary statistic for the reliability maps. We used bootstrap tests to estimate confidence intervals and statistical significance. In each of 20,000 repetitions, we randomly sampled a group of 20 individuals by sampling with replacement from the 20 original individuals, then computed IDMs based on the bootstrapped sample, and obtained statistics accordingly, such as average reliability and average reliability difference. During bootstrapping, non-informative correlation values originally in the diagonal of an IDM (self-similarity values of 1 for a given participant due to sampling with replacement) may appear in off-diagonal cells, so we excluded these values from the upper triangles to avoid overestimation of reliability values (Kriegeskorte et al., 2008).

### 2.7 Indexing local functional architecture

We measured individual differences in local functional architecture based on three functional indices that are commonly used in the neuroimaging literature: response profiles, functional connectivity (Biswal et al., 1995), and representational geometry (Kriegeskorte and Kievit, 2013; Kriegeskorte et al., 2008).

We use the term “response profile” to refer to the response time series to a part of the movie for each feature in searchlight. The response profiles of all features in the searchlight formed a time-points by features data matrix for each individual. This data matrix was vectorized into a vector of response profiles reflecting the spatiotemporal pattern of an individual’s responses to the movie. We constructed response profile IDMs by computing the similarity of these response profile vectors across all pairs of individuals. We computed the functional connectivity profile of each feature as the Pearson correlation between its response time series and the response time series for all 18,742 features in the brain, which reflects its co-activation pattern with those 18,742 connectivity targets. The functional connectivity profiles of all features in a searchlight formed a connectivity targets by features matrix, which can be vectorized and compared across individuals to form functional connectivity IDMs. We computed the representational geometry of each searchlight as a time-point-based representational dissimilarity matrix (RDM) (Guntupalli et al., 2016; Kriegeskorte et al., 2008) based on correlation distance. That is, for each searchlight we first computed a time-points by time-points RDM. Then, the similarity between each pair of individuals was measured as the correlation between vectorized upper triangles of the RDMs, resulting in representational geometry IDMs.

### 2.8 Separating coarse-scale and fine-scale individual differences

We divided the information in those functional indices into coarse-scale information and fine-scale information, to test if the alignment method used affects individual differences differently based on the spatial scale of topographic functional information.

When using response profiles as the functional index, we defined coarse-scale information as the mean response profile (time series) across all features in a searchlight, and fine-scale information as the pattern residuals after subtracting the searchlight mean response profile from each feature’s response profile. In this case, the remaining pattern only contains fine-scale information and no variation of mean response across movie time-points. When using functional connectivity as the functional index, we defined coarse-scale information as the correlations between the searchlight mean response profile and those from the 18,742 connectivity targets, and the fine-scale information as the correlations between the residual time series of the features and those of the targets. We measured representational geometry as correlation distances between spatial patterns of different movie time-points. Correlations between spatial patterns implicitly remove the mean and standard deviation across features (Misaki et al., 2010), and thus only use fine-scale information. Therefore, we didn’t perform this analysis with representational geometry.

### 2.9 Comparing functional indices

We then compared IDMs based on different functional indices to see if the structure among individuals based on one functional index was retained for the other functional indices. Thus, instead of comparing IDMs from different parts of the movie data based on the same functional index (as in the reliability analysis, see 2.6; e.g., index 1 movie part 1 vs. index 1 movie part 2), we compared IDMs from different parts of the movie based on different functional indices. In this approach, two correlation coefficients can be obtained from each pair of functional indices (index 1 movie part 1 vs. index 2 movie part 2, and index 1 movie part 2 vs. index 2 movie part 1). We averaged these two correlation coefficients to reduce estimation error.

## 3. Results

### 3.1 Reliability of individual differences

We computed searchlight maps of the reliability of individual differences in response profiles, functional connectivity, and representational geometry using both anatomically aligned data and hyperaligned data. Each location on the cortical surface is assigned a correlation value (ranging from -1 to 1) denoting the reliability of individual differences for a given functional index in that searchlight. Note that we expect most reliabilities to be positive, and that negative and near-zero reliabilities are indicative of noise.

The average reliability of individual differences based on response profiles across all searchlights was 0.540 for anatomically aligned data (95% CI of [0.446, 0.585] estimated by bootstrapping subjects); the average reliability was 0.693 [0.578, 0.744] for hyperaligned data (Fig. 1B and 1C). Mean reliability was significantly higher for hyperaligned data than for anatomically aligned data, with an average increase in reliability of 0.153 [0.116, 0.182], *p* < 10^-4^. Most searchlights (82.1%) had higher reliability with hyperalignment, consistent with the searchlight average results, indicating that the increase in average reliability was not due to outlier searchlights.

The average reliability of individual differences based on functional connectivity (Fig. 2, upper) across all searchlights increased from 0.799 [0.750, 0.827] for anatomically aligned data to 0.861 [0.808, 0.894] for hyperaligned data, an average increase in reliability of 0.062 [0.020, 0.106], *p* = 0.008. Across all searchlights, 72.7% had higher reliability with hyperalignment.

**Fig. 2.**
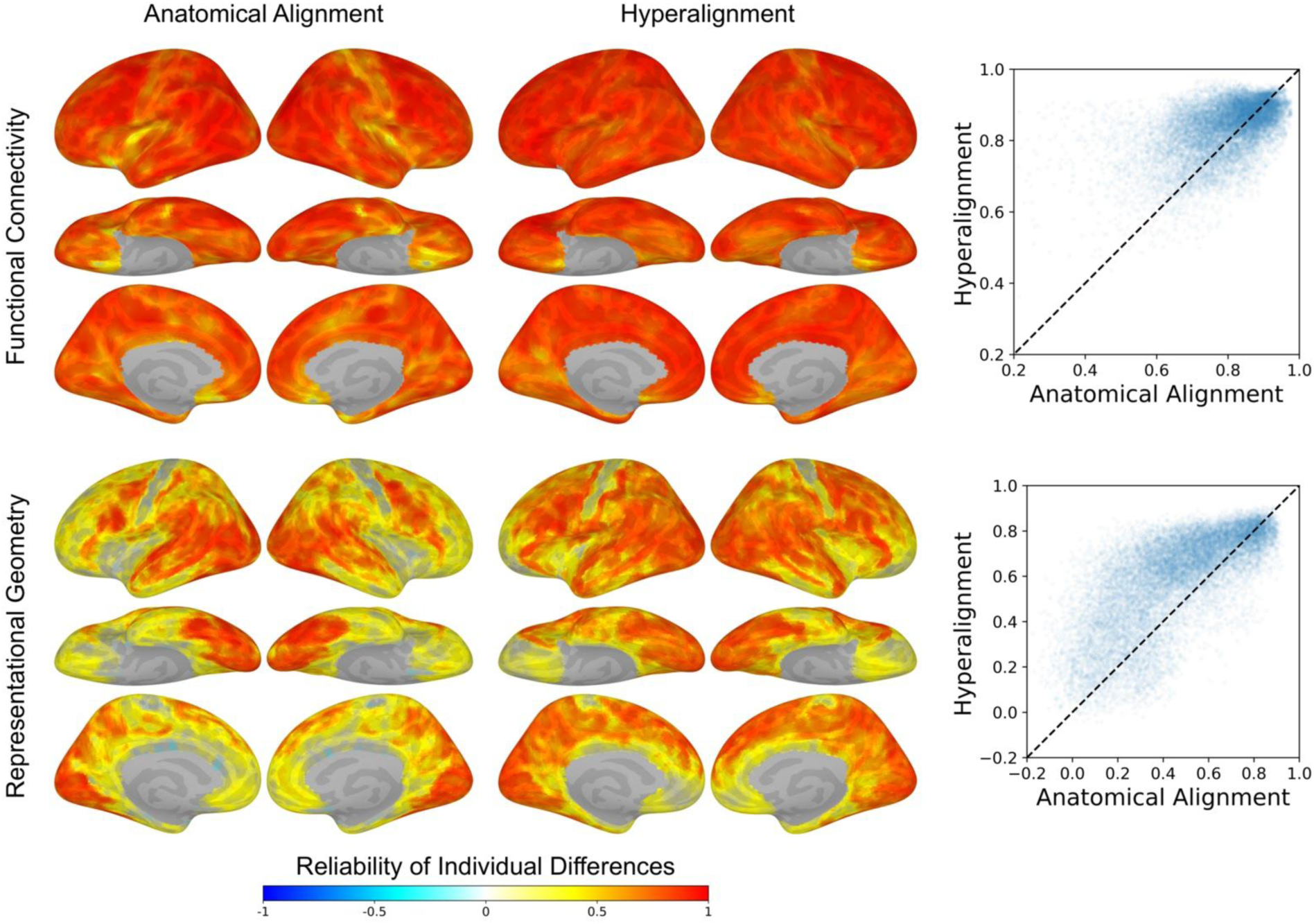
The reliability of individual differences indexed by functional connectivity (upper) and representational geometry (lower). When these alternative functional indices were used, individual differences also became more reliable with hyperalignment. When using functional connectivity to index individual difference, 72.7% searchlights had higher reliability with hyperalignment, and the average reliability increased from 0.799 to 0.861. When using representational geometry, 77.5% had higher reliability with hyperalignment, and the average reliability increased from 0.472 to 0.601.

The average reliability of individual differences based on representational geometry (Fig. 2, lower) across all searchlights increased from 0.472 [0.387, 0.499] with anatomically aligned data to 0.601 [0.482, 0.657] with hyperaligned data, an average increase in reliability of 0.129 [0.089, 0.166], *p* < 10^-4^. Across all searchlights, 77.5% had higher reliability with hyperalignment.

In summary, the reliability of individual differences in response profiles, functional connectivity, and representational geometry all became more reliable after hyperalignment. Across the three functional indices, individual differences in functional connectivity were more reliable than individual differences in response profiles, and individual differences in response profiles were more reliable than individual differences in representational geometry. Based on anatomically aligned data, 91.2% searchlights had higher reliability for functional connectivity compared with response profiles, and the average difference across all searchlights was 0.260 [0.205, 0.344]; 65.5% searchlights had higher reliability for response profiles compared with representational geometry, and the average difference across searchlights was 0.068 [0.043, 0.103]. Based on hyperaligned data, 92.8% searchlights had higher reliability for functional connectivity compared with response profiles, and the average difference across all searchlights was 0.168 [0.110, 0.267]; 75.6% searchlights had higher reliability for response profiles compared with representational geometry, and the average difference across searchlights was 0.092 [0.067, 0.126].

### 3.2 Coarse-scale and fine-scale individual differences

We separated the information encoded in each searchlight into coarse and fine spatial scales, and assessed the reliability of individual differences for both. Coarse-scale information is defined as the average time series across all cortical features within a searchlight (regional-average response profile) and the correlations of the searchlight average time series with functional connectivity targets (regional-average connectivity profile), and the fine-scale information is based on the residuals of the data matrix after removing the searchlight average time series or average functional connectivity vector. These approaches characterized independent and complementary types of information encoded in each searchlight.

When using response profiles to index individual differences (Fig. 3, upper), the average reliability based on coarse-scale information marginally increased from 0.496 [0.400, 0.542] with anatomically aligned data to 0.528 [0.411, 0.590] with hyperaligned data, an average reliability difference of 0.032 [0.000, 0.059], *p* = 0.049. Across all searchlights, 58.8% had higher reliability with hyperalignment. By contrast, the average reliability based on fine-scale information increased from 0.415 [0.348, 0.455] with anatomically aligned data to 0.660 [0.538, 0.714] with hyperaligned data, an average increase in reliability of 0.246 [0.181, 0.282], *p* < 10^-4^. Across all searchlights, 92.5% had higher reliability with hyperalignment.

**Fig. 3.**
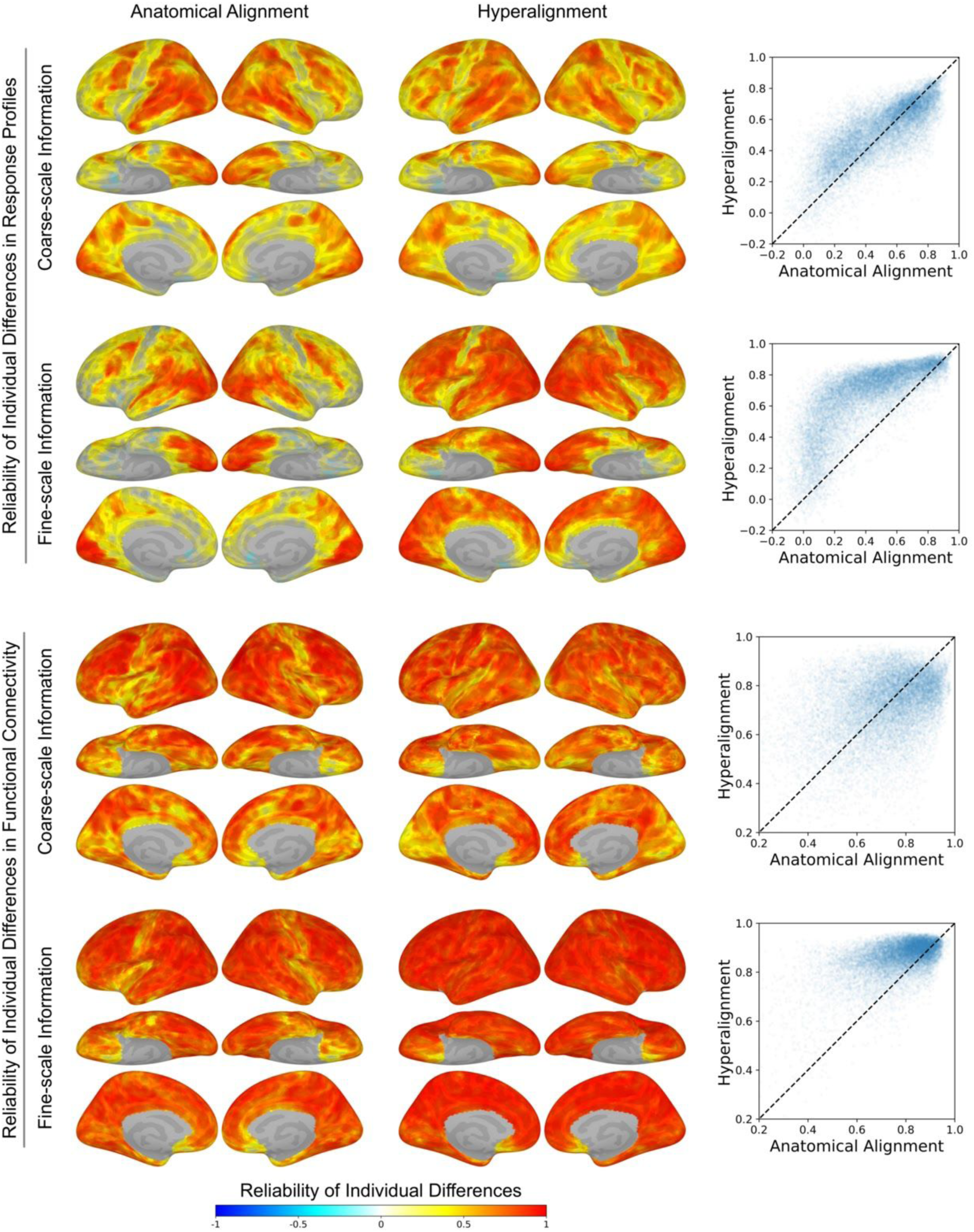
Individual differences at coarse and fine spatial scales. We separated coarse-scale information (regional-average response profiles or regional-average connectivity profiles) and fine-scale information (pattern residuals after removing coarse-scale information) from the functional indices, and modeled individual differences accordingly. When using response profiles to index individual differences, hyperalignment increased the average reliability of coarse-scale individual differences from 0.496 to 0.528, and that of fine-scale individual differences from 0.415 to 0.660. When using functional connectivity, hyperalignment increased the average reliability of coarse-scale individual differences from 0.726 to 0.730, and that of fine-scale individual differences from 0.782 to 0.865. In general, the reliability of fine-scale individual differences benefited more from hyperalignment compared with coarse-scale individual differences.

When using functional connectivity to index individual differences (Fig. 3, lower), the average reliability based on coarse-scale information was similar for anatomically aligned data (0.726 [0.679, 0.749]) and hyperaligned data (0.730 [0.673, 0.772]), with an average reliability difference was 0.004 [-0.044, 0.057], *p* = 0.796. Across all searchlights, 48.2% had higher reliability with hyperalignment. The average reliability based on fine-scale information increased from 0.782 [.0754, 0.792] with anatomically aligned data to 0.865 [0.810, 0.898] with hyperaligned data, an average increase in reliability of 0.083 [0.040, 0.121], *p* = 0.003. Among all searchlights, 81.3% had higher reliability with hyperalignment.

The reliability of fine-scale individual differences benefited more from hyperalignment than did the reliability of coarse-scale individual differences (Fig. 4). When using response profiles to index individual differences, the increases in average reliability with hyperalignment were 0.246 and 0.032 respectively, and the increase for fine-scale individual differences was higher by 0.213 [0.165, 0.239], *p* < 10^-4^. Hyperalignment yielded a larger increase in the reliability of fine-scale individual differences in 88.9% of searchlights. When using functional connectivity to index individual differences, the increases in average reliability were 0.083 and 0.004 respectively, and the increase for fine-scale individual differences was higher by 0.079 [0.042, 0.117], *p* < 10^-4^. Across all searchlights, 71.4% had a larger increase in the reliability of fine-scale individual differences. Note with both functional indices, fine-scale individual differences based on hyperaligned data had the highest average reliability among the four combinations, thus the smaller increase in the reliability of coarse-scale individual differences was not due to ceiling effects.

**Fig. 4.**
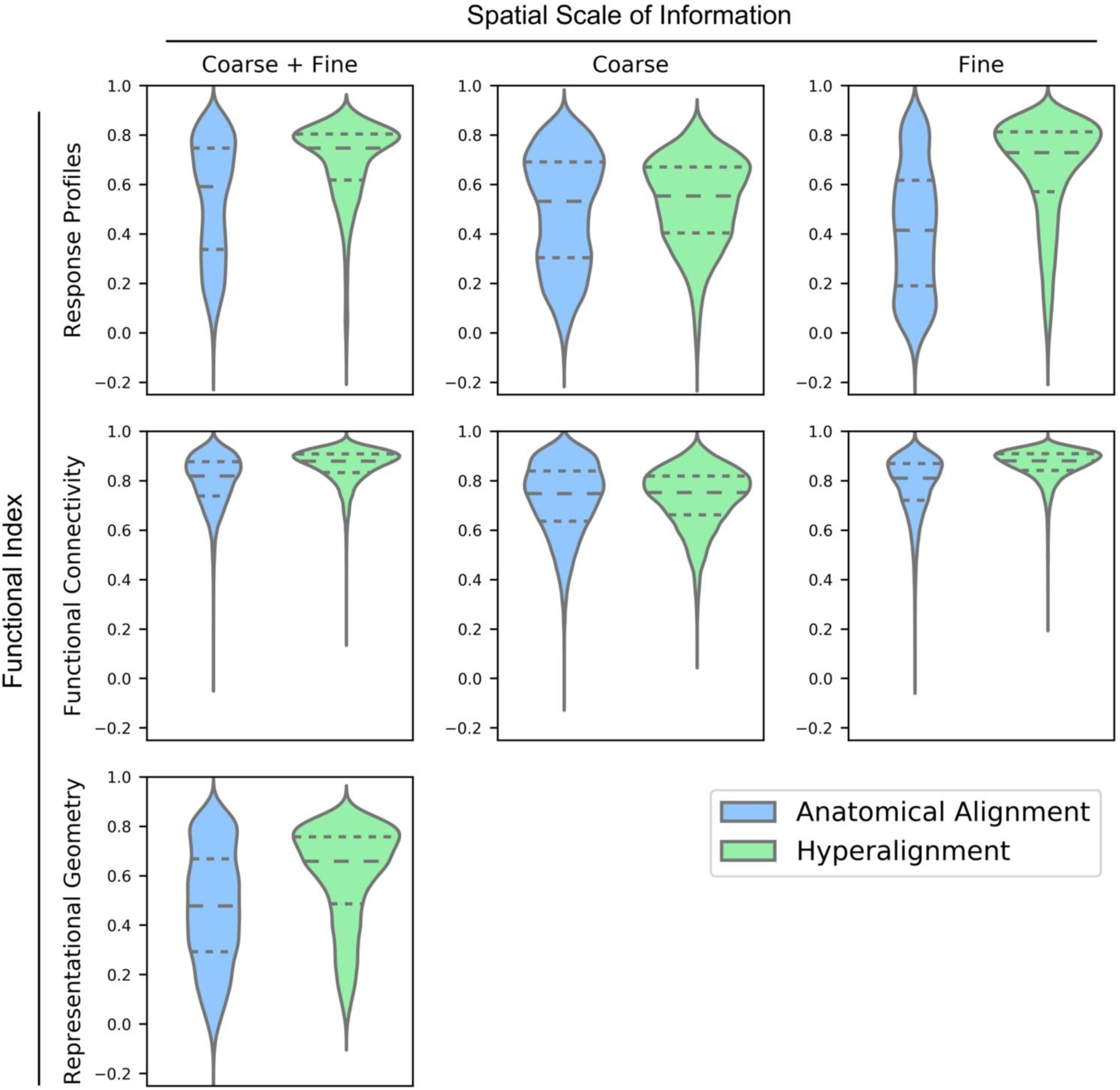
Distribution of searchlight reliabilities of individual differences across functional indices, spatial scales, and alignment methods. With hyperalignment, individual differences were more reliable across all three functional indices (left column, see also Fig. 1B and Fig. 2). When only coarse-scale information was used, the distribution of reliabilities was similar for both alignment methods (middle column, see also Fig. 3). By contrast, when only fine-scale information was used, the distribution of reliabilities shifted toward higher values with hyperalignment (right column, see also Fig. 3). Dashed lines denote quartiles of each distribution.

### 3.3 Agreement across functional indices

We next correlated IDMs based on different functional indices derived from different parts of the movie data (Fig. 5). A positive correlation indicates that individuals who differ more according to one functional index are also likely to differ more according to the other functional index, and individuals who are more similar on one index are also more similar on the other index. Inter-index correlations computed across the two parts of the movie were then averaged. Consequently, the similarity of IDMs based on different indices is derived from different parts of the movie for each index.

**Fig. 5.**
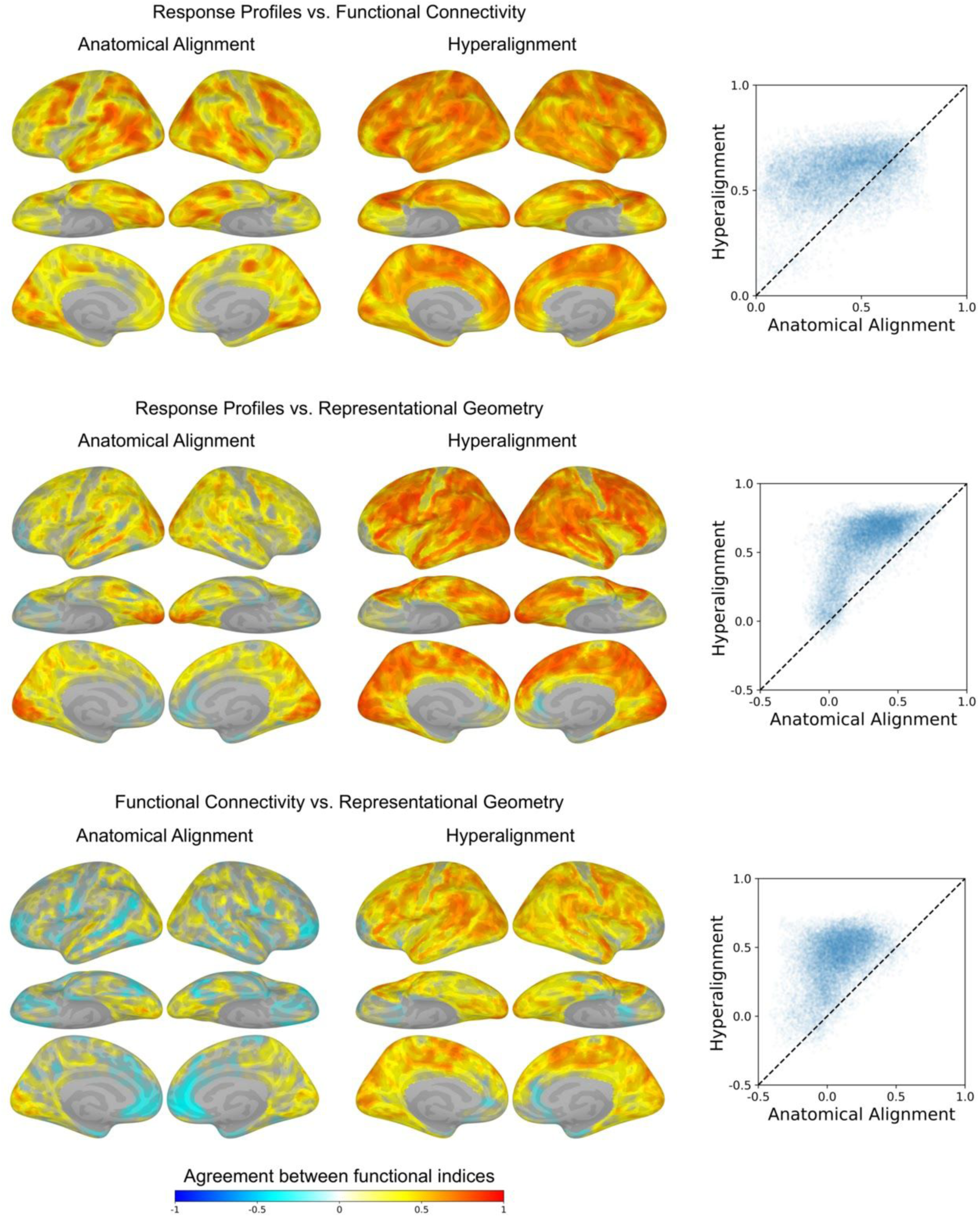
Agreement of individual differences across functional indices. Within each searchlight, we computed the correlation between two IDMs based on different functional indices and different parts of the movie data, and averaged that across permutations of movie parts. A positive correlation means that individuals who differ more according to one functional index are likely to differ more for the other functional index. The average correlation between response profile IDMs and functional connectivity IDMs was 0.408 for anatomically aligned data, and 0.582 for hyperaligned data. The average correlation between response profile IDMs and representational geometry IDMs was 0.272 for anatomically aligned data, and 0.554 for hyperaligned data. The average correlation between functional connectivity IDMs and representational geometry IDMs was 0.077 for anatomically aligned data, and 0.415 for hyperaligned data. After hyperalignment, individuals who differed along one functional index were more likely to differ for others.

When comparing individual differences in response profiles and individual differences in functional connectivity, with anatomically aligned data, 99.5% of searchlights had positive correlations, and the average correlation across all searchlights was 0.408 [0.334, 0.465]; with hyperaligned data, 100.0% of searchlights had positive correlations, and the average correlation across all searchlights was 0.582 [0.454, 0.668]. Among all searchlights, 83.8% had higher correlations with hyperaligned data, and hyperalignment increased the average correlation by 0.174 [0.093, 0.247], *p* < 10^-4^.

When comparing individual differences in response profiles and individual differences in representational geometry, with anatomically aligned data, 91.3% of searchlights had positive correlations, and the average correlation across all searchlights was 0.272 [0.203, 0.317]; with hyperaligned data, 98.4% of searchlights had positive correlations, and the average correlation across all searchlights was 0.554 [0.422, 0.626]. Among all searchlights, 96.5% had higher correlations with hyperaligned data, and hyperalignment increased the average correlation by 0.281 [0.216, 0.315], *p* < 10^-4^.

When comparing individual differences in functional connectivity and individual differences in representational geometry, with anatomically aligned data, 66.4% of searchlights had positive correlations, and the average correlation across all searchlights was 0.077 [0.018, 0.135]; with hyperaligned data, 96.0% of searchlights had positive correlations, and the average correlation across all searchlights was 0.415 [0.263, 0.520]. Among all searchlights, 96.8% had higher correlations with hyperaligned data, and hyperalignment increased the average correlation by 0.338 [0.229, 0.417], *p* < 10-4.

Compared with anatomically aligned data, more searchlights had positive inter-index IDM correlations with hyperaligned data, and the average correlation across all searchlights increased. Thus, after hyperalignment, all three functional indices converged on more congruent characterizations of inter-subject similarity of cortical functional architecture.

## 4. Discussion

In this study, we measured individual differences in local cortical functional architecture based on three functional indices—response profiles, functional connectivity, and representational geometry-using dynamic, naturalistic movie stimuli. We quantified individual differences with IDMs. Each IDM provides an index of the similarity structure among individuals, which can be thought of as an individual-differences geometry. IDMs derived from different parts of the movie or from different functional indices can be compared directly. These IDMs capture neural similarities among subjects. The relationship of these individual differences in cortical functional architecture to phenotypic variation is beyond the scope of this study—however, reliable neural IDMs can be considered a precondition for relating individual differences in neural function to these other variables (Castellanos et al., 2013; Dubois and Adolphs, 2016; Gabrieli et al., 2015; Van Horn et al., 2008; Woo et al., 2017). We assessed the reliability of individual differences by comparing IDMs from different parts of the movie, and found reliable individual differences in all three functional indices throughout cortex. Naturalistic stimuli are engaging and content-rich, evoking a variety of brain states and facilitating the detection of individual differences (Vanderwal et al., 2017). Importantly, after aligning each individual’s data to a common space using hyperalignment, we found individual differences in all three functional indices become even more reliable; hyperalignment increased reliability in a clear majority of searchlights (73% to 82%) and increased average reliability across all searchlights. This suggests individuals differ reliably in local cortical functional architecture, but that these differences can be obscured by suboptimal alignment of brain imaging data across individuals.

To further test this hypothesis, we divided the information in these functional indices into coarse-scale information and fine-scale information, and assessed the reliability of individual differences accordingly. Coarse-scale information was computed based on the average time series of all nodes in a searchlight, and thus was spatially smooth and relatively robust against misalignment of data across individuals (conceptually related to the motivation for spatially smoothing functional data in multi-subject univariate analyses; Petersson et al., 1999; Carp, 2012). The information not captured by searchlight average time series, i.e., the residual time series in each cortical vertex, was defined as fine-scale information, and thus was independent and orthogonal to coarse-scale information. We found the reliability of coarse-scale individual differences was the same or slightly increased with hyperalignment. In contrast, the reliability of fine-scale individual differences increased dramatically with hyperalignment, and surpassed the reliability of coarse-scale individual differences. These results suggest individual differences in local cortical functional architecture exist at both coarse and fine spatial scales. However, individual differences in fine-scale functional architecture can only be efficiently captured when data are properly aligned across individuals. This is a critical development for identifying biomarkers, as some abnormalities may not be evident in coarse-grained functional organization and only manifest at fine spatial scales (Woo et al., 2017; cf. Hackmack et al., 2012).

It is essential to establish functional correspondence across individuals before analyzing how they differ in cortical functional architecture. Consider measuring a single brain feature (e.g., voxel, vertex, electrode, etc.) at the same macroanatomical position in two individuals that responds preferentially to different kinds of stimuli. The two individuals’ response time series would be highly similar if the two kinds of stimuli always co-occur, and highly dissimilar if the two kinds of stimuli always appear in an interleaved fashion. In these scenarios, measured individual differences in cortical functional architecture are prone to the co-occurrence and frequency of stimuli, and less generalizable to new tasks and new sets of stimuli. On the other hand, if the same brain feature from the two individuals are from a common functional space (e.g., common space created by hyperalignment), they are expected to share similar functional properties like response tuning profiles or functional connectivity profiles. In this case, measured individual differences in cortical functional architecture will be robust against variations in stimuli, and thus more generalizable and reliable.

Consider the same brain feature with differential response tuning across two individuals. Is this discrepancy due to topographic differences? The mismatch may be due to differences in functional-anatomical correspondence across individuals, or the way in which a particular subject’s brain signals were sampled (e.g., the discretization of brain signals into voxels). In this simplistic example, the discrepancy may be resolved simply by remapping one feature to another nearby feature across individuals. Hyperalignment, and related approaches (e.g., Yamada et al., 2015), relax this one-to-one mapping and flexibly account for topographic differences by modeling the responses of a given feature as a weighted sum of nearby features in the common space. If, after hyperalignment, the discrepancy remains, this is strongly indicative of a feature in a particular individual that deviates from the group model for the cortical field to which that feature belongs. In a multivariate context, this would be reflected in individual differences in representational geometry (Charest et al., 2014). By accounting for idiosyncrasies in functional-anatomical correspondence, hyperalignment offers a rigorous framework for modeling deviations from the group and a less-confounded view of individual differences in function.

At first glance, one might expect that hyperalignment would effectively eliminate individual differences. Hyperalignment does in fact increase similarities among individuals for the functional indices we use (Guntupalli et al., 2016, in press; Haxby et al., 2011), but nonetheless preserves (and enhances) the structure of the similarities among individuals. The hyperalignment implementation used in this paper uses Procrustes transformations within searchlights, which is a rotation in searchlight high-dimensional feature spaces. It re-distributes variances across features rather than changing the variances themselves. Therefore, the Procrustes transformation is expected to correct for individual differences in topographic distributions of function while maintaining individual differences in function per se. For example, if one individual’s response magnitude to a stimulus is twice as high as another individual, or if the response exists in twice as many features, such differences will still be retained in Procrustes-transformed data. However, if two individuals have the same response but the response exists in different features, their differences will be resolved by the transformation. Therefore, by projecting each individual’s data into the common space, we can measure differences in functional tuning without confounds from (mis)localization of function.

Searchlight hyperalignment (Guntupalli et al., 2016) adds local Procrustes transformations together to form a whole-brain transformation matrix. When a feature is included in multiple searchlights, respective transformations are combined. This procedure is related to ensemble learning techniques (Zhou, 2012) in that it boosts the shared part of the transformations and suppresses the noisy part. Therefore, instead of a completely orthonormal whole-brain transformation matrix, it provides a more accurate and robust whole-brain transformation. As a result, the signal shared across individuals will be boosted and the noise suppressed; this may partly explain why individual differences in cortical functional architecture become even more reliable after searchlight hyperalignment.

Representational geometry (Kriegeskorte and Kievit, 2013) measures the structure of similarities between stimulus representations, which itself is not expected to change with rotations in the feature space (e.g., the Procrustes transformation). However, variances in a searchlight after hyperalignment are not identical to the variances in the searchlight in anatomically aligned data. Small differences in representational geometry may be observed after searchlight hyperalignment partly because the method for aggregating searchlights will adjust which (and to what extent) features are considered “members” of a given cortical field. This is conceptually analogous to localizing functional regions of interest (ROIs) in univariate analyses (Saxe et al., 2006), and comparing these functional ROIs rather than comparing anatomical ROIs (which increases statistical power and functional resolution; Nieto-Castañón and Fedorenko, 2012). Similarly to functional ROIs, following hyperalignment, representational geometry is computed across individuals in a “functional searchlight” instead of a purely anatomy-based searchlight. However, note the “boundary” of the “functional searchlight” is not an anatomically-defined boundary on surface, but is rather a boundary in a high-dimensional information space determined by a transformation which maps multivariate information from each individual’s anatomical space to the information searchlights in common space.

Therefore, measuring individual differences in representational geometry with searchlight hyperalignment also benefits from better alignment of functional architecture and reduced topographic confounds.

Although individual differences in local cortical functional architecture can be based on various functional indices, we found that the structure of individual differences was similar across the three functional indices, especially following hyperalignment. Note that individual differences in functional topography affect these functional indices to different degrees: representational geometry is designed to be independent of within-area topographic differences (rotations in the feature space); response profiles can be affected by topographic differences to some degree; functional connectivity is most prone to topographic differences, as both features in an area and connectivity targets can be affected by topographic differences. Such effects are expected to be reduced by hyperalignment, and the individual differences measured in the hyperaligned common space mainly reflect functional differences without confounding topographic differences. Each functional index is differentially susceptible to topographic differences and hyperalignment improves agreement across indices by attenuating these topographic differences.

Hyperalignment yields more reliable measures of individual differences in cortical functional architecture both by reducing confounds from topographic idiosyncrasies and by capturing variation around shared bases for how information is encoded in fine-scale topographic patterns. This is a promising step forward for efforts to link individual differences in brain function to individual differences in behavior. As translational neuroscience matures (Dubois and Adolphs, 2016; Gabrieli et al., 2015; Poldrack, 2017; Woo et al., 2017), hyperalignment will be instrumental in building more detailed, clinically-relevant biomarkers.

## Acknowledgements

We thank Yaroslav O. Halchenko, M. Ida Gobbini, Andrew C. Connolly, Matteo Visconti di Oleggio Castello, and Vassiki Chauhan for helpful discussions.

